# Glial cell-intrinsic and non-cell autonomous toxicity in a Drosophila C9orf72 neurodegeneration model

**DOI:** 10.1101/2025.10.07.680869

**Authors:** I. Hubbard, J. Dubnau

## Abstract

The most common genetic cause of both familial amyotrophic lateral sclerosis (ALS) and frontotemporal dementia (FTD) is an expanded G_4_C_2_ repeat in the first intron of the gene C9orf72. The C9orf72 repeat expansion is bidirectionally transcribed into sense and anti-sense RNA foci, and also produces dipeptide repeats (DPRs) via a non-canonical translation mechanism known as repeat-associated (RAN) translation. Each of these components of the G_4_C_2_ repeat expansion cause neurodegenerative effects in animal models when expressed in neurons, but impacts from glial expression are more poorly understood. Here, we use glial cell type-specific expression of individual DPRs, of RNA repeat-only, or of the G_4_C_2_ repeat that is capable of producing both DPRs and RNA repeats to systematically investigate both the glial cell-intrinsic and non-cell autonomous toxicity of each of these components. Our results show that as with neurons, the GR and G_4_C_2_ transgenes, produce the highest degree of cell-intrinsic toxicity when expressed in glia. Both of these transgenes are capable of producing the GR DPR, which is also typically found to be the most toxic factor in neurons. We demonstrate that both the GR and G_4_C_2_ transgenes cause activation of mdg4, an endogenous retrovirus (ERV). Such ERV expression is a hallmark of TDP-43 dysfunction that is commonly observed in C9orf72 patients and contributes to both cell intrinsic and non-cell autonomous toxicity. We find that only the G_4_C_2_ transgene produces measurable non-cell autonomous effects that result in loss of nearby neurons. But manipulations of apoptosis reveal non-cell autonomous or systemic effects from either GR or G_4_C_2_ expressing glia. Blocking apoptotic cell death of either GR or G_4_C_2_ expressing glia via the p35 caspase inhibitor further exacerbates effects on lifespan and ablating such glia via expression of the proapoptotic reaper gene partially ameliorates these effects.

## Introduction

Amyotrophic lateral sclerosis (ALS) and frontotemporal dementia (FTD) are two fatal neurodegenerative diseases that exist on a clinical spectrum(1). On a biological level, ALS and FTD feature shared proteinopathy, with over 90% of ALS patients and approximately 50% of FTD patients exhibiting abnormal cytoplasmic aggregation of TAR-DNA binding protein 43 (TDP-43) (2). Although most cases have no known genetic cause, a subset of cases are familial. The most common genetic form of both ALS and FTD is a hexanucleotide GGGGCC (G_4_C_2_) repeat expansion in the gene C9orf72, which can exist in up to hundreds or even thousands of repeats in patients (3–5). The pathological impacts of this repeat expansion are complex. First, there is a potential toxic loss of function, as blood lymphocyte and tissue samples from C9orf72 expansion carriers exhibit reduced C9orf72 encoded transcript levels compared to controls (3,4,6,7). However, multiple studies have failed to demonstrate motor and behavioral deficits in knockout models, suggesting that loss-of-function by itself is insufficient to drive ALS/FTD disease (8). The structure of the repeat expansion is also capable of producing two putative toxic gain-of-function mechanisms. First, the repeat can be transcribed in both the sense and antisense directions to produce RNA foci, which accumulate both inside and outside the nucleus (9,10). Additionally, the repeat is capable of being translated from both strands and in all reading frames via a non-canonical translation mechanism known as repeat associated non-AUG (RAN) translation. RAN translation of both the sense and anti-sense transcripts can give rise to five different dipeptide repeats (DPRs): glycine-proline (GP), glycine-alanine (GA), glycine-arginine (GR), proline-alanine (PA), and proline-arginine (PR). GA and GR are produced by the sense strand, PA and PR are produced by the antisense strand, and GP is produced by both strands(9,11,12). In addition to this complex mixture of pathological hallmarks that involve products of the repeat expansion in the *C9orf72* gene, TDP-43 pathology is also observed in C9orf72 familial cases (1). Thus, elucidating the contributions of each pathological feature to C9orf72*-*related ALS and FTD is a considerable challenge.

Numerous studies have examined the differential toxic effects of RNA foci and DPRs when expressed in neurons, with the arginine-rich DPRs GR and PR typically exhibiting the greatest degree of toxicity (13–15). Considerable effort has gone to investigate the mechanisms by which these arginine rich DPRs cause cell intrinsic toxicity to neurons (16–18). But less is known about the potentially synergistic impacts of expressing combinations of DPRs, about their toxicity to glial cells, and about the non-cell autonomous effects of expressing each of these components that are encoded by this G_4_C_2_ expansion repeat.

Non-cell autonomous effects are a key feature of neurodegenerative disease. In the case of both ALS and FTD, there is strong evidence for progressive spread of disease pathology from a focal origin of onset (19–21). Such non-cell autonomous toxicity also involves signaling between several different cell types, including neurons and glial cells. In ALS and FTD, astrocytes are thought to release toxic substances with negative impact on motor neuron survival. This phenomenon of glial-to-neuronal toxicity has been shown *in vivo* in a transgenic rat model in which mutant TDP-43 expression is restricted to astrocytes (22), as well as in primary astrocyte-motor neuron co-culture, including from *C9orf72* patient-derived iPSC astrocytes, which confer toxicity to motor neurons in co-culture and organoid models (23–25). Similar non-cell autonomous toxicity from glia to neurons has been observed in *Drosophila* models of TDP-43 pathology (26,27), but have not been systematically studied in the context of *C9orf72* [but see(28)].

We have previously demonstrated that initiating TDP-43 protein pathology via overexpression in *Drosophila* glia causes motor defects and reduction in lifespan (29). Such glial over-expression of TDP-43 also causes DNA damage and cell death in nearby neurons (26). In particular, focal over-expression of human TDP-43 to drive protein pathology within a specialized glial subtype, the subperineural glia (SPG), is sufficient to cause appearance of TDP-43 protein pathology and toxicity in nearby neurons that express physiological levels of TDP-43. This non-cell autonomous effect is mediated at least in part by de-repression in the SPG of mdg4, an endogenous retrovirus (ERV) (27). ERVs and other retrotransposable elements (RTEs) are aberrantly over-expressed in *Drosophila* models, in a mouse model, and also in postmortem cortical tissues from human ALS/FTD patients and in human neuroblastoma cells that exhibit TDP-43 protein pathology (26,27,29–34). Here, we investigated both the cell intrinsic and non-cell autonomous effects of expressing C9orf72 repeats in *Drosophila* glia. We compared the effects of pan glial and cell type-specific expression of individual DPRs, of RNA repeat-only, or of the G_4_C_2_ repeat that is capable of producing both DPRs and RNA repeats.

## Results

### Pan-glial expression of arginine-rich dipeptide repeats drives toxicity that shortens lifespan

We have previously demonstrated that pan-glial over expression of human TDP-43 in *Drosophila* reduces lifespan and causes non-cell autonomous toxicity to nearby neurons(26). Such over-expression triggers clearance from the nucleus and appearance of cytoplasmic inclusions of hyperphosphorylated TDP43. Since TDP-43 protein pathology is a feature that is observed in postmortem tissue of C9orf72 cases (35), we sought to examine whether expression of individual components of the C9orf72 repeat expansion in glia might recapitulate the toxicity observed with glial overexpression of TDP-43. In order to achieve post-development glial expression, we first used a pan-glial Gal4 line (Repo-Gal4) in combination with a temperature sensitive Gal80 (Gal-80^ts^). Using this Gal80^ts^ method, we induced expression of a series of previously reported (13) UAS-transgenes that each express different subsets of the components that could contribute to toxicity of the G_4_C_2_ repeat expansion. In each case, we kept the length of the repeat constant (at 36 repeats). These transgenes consisted of a (G_4_C_2_)_36_ repeat capable of producing both the repeat RNA and the sense DPRs, an RNA-only (G_4_C_2_)_36_ RO repeat that is unable to produce any of the DPRs, and protein-only constructs that do not make an RNA repeat but are capable of producing individual dipeptide repeat proteins. For this last category of DPR only transgenes, we used transgenes that produce individual sense-derived DPRs-glycine-alanine (GA_36_) and glycine-arginine (GR_36_), or the antisense DPRs-proline-alanine (PA_36_), and proline-arginine (PR_36_) (13).

For each transgene, we used the Gal80^ts^ approach to induce glial expression of the above transgenes after development was completed. This approach allowed us to avoid the potential effects from expressing these various C9orf72 repeat-associated pathological proteins or RNAs during development. We first examined the effects on lifespan from pan-glial expression of each of the C9 UAS transgenes (Figure 1A). The G_4_C_2_ repeat and sense-derived GR-expressing DPR construct produced the most detrimental effects on lifespan. The other arginine-rich DPR, the antisense-derived PR, also appeared to have a robust effect on lifespan, although the effects were not as severe as with the G_4_C_2_ or GR transgenes. Expression of the RNA-only repeat yielded a significant impact on lifespan, although to a lesser degree than the G_4_C_2_ or GR DPR. The PA-expressing DPR had little impact, and the GA-expressing DPR had no significant impact. Examining lifespan effects of each of these DPRs of length 100 shifted median survival only 1-2 days compared to their counterparts with 36 length repeats, indicating that at least for such glial expression, repeat content is more important than length in promoting toxicity (Figure S1). Taken together, these findings are consistent with the interpretation that the ability to produce the arginine-rich GR DPR is a driving contributor to the toxicity produced by the sense G_4_C_2_ repeat.

**Figure 1.**
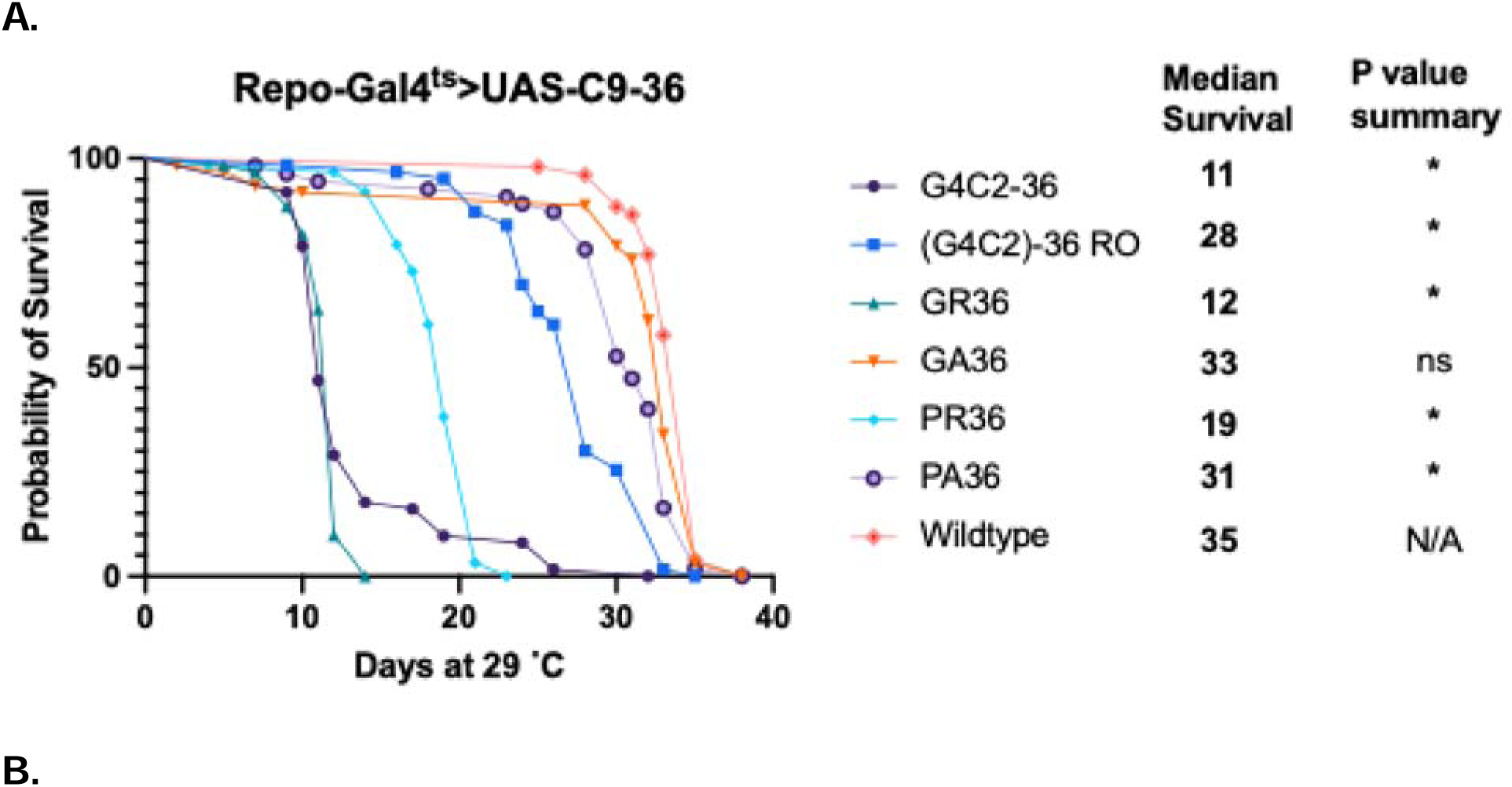
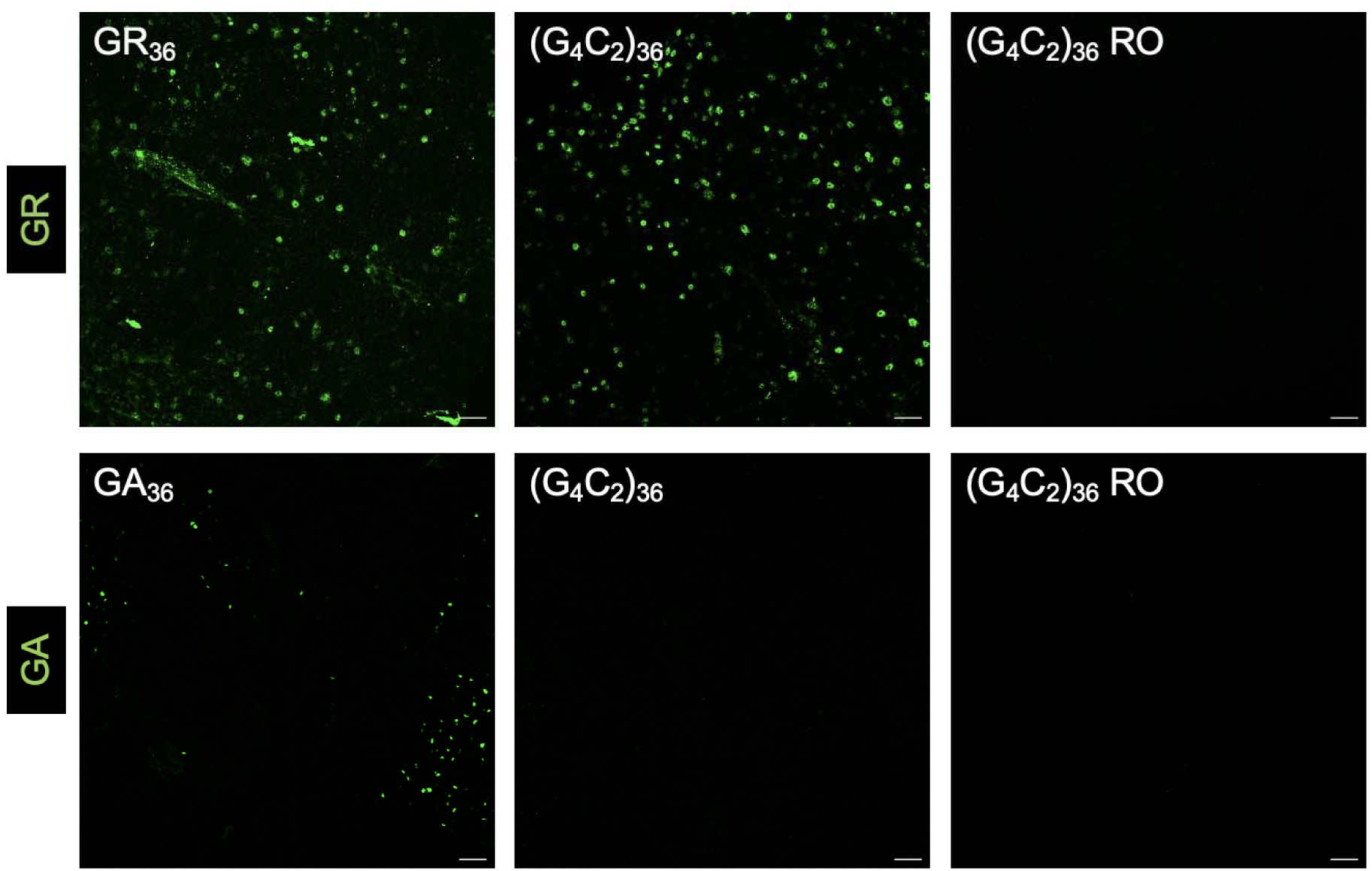
Pan-glial expression of arginine-rich repeats drives toxicity that shortens lifespan. (A). Lifespan analysis of flies with pan-glial expression of C9 UAS transgenes, compared to a wildtype control containing the Repo-Gal4^ts^ driver. *p < 0..00238 (log-rank test with Bonferroni’s correction) (B). Confocal images of *Drosophila* brains at 40x magnification stained with anti-GR (top row) or anti-GA (bottom row) antibodies after 3 days of pan-glial expression. Both the GR and GA UAS transgenes (left column) expressed their respective DPR. (G_4_C_2_)_36_ (middle column) expressed abundant GR; GA was not detected. Neither GR nor GA was detected in (G_4_C_2_)_36_ RO (right column). Scale bar = 20 µm. *Full genotype: Repo-Gal4, Tubulin-Gal80^ts^ >UAS-C9*

We next sought to verify that the G_4_C_2_ repeat in fact produced the expected DPRs in glia. Since it has previously been reported that the G_4_C_2_ model does not produce antisense transcripts (13), we specifically examined the sense DPRs GR and GA. Immunostaining using antibodies against GR and GA three days after induction of each respective C9orf72-related transgene revealed that GR was in fact abundantly expressed from both the G_4_C_2_ and the GR specific transgenes. By contrast, we are unable to detect GA expression from the G_4_C_2_ repeat transgene, although we do detect accumulation of the GA DPR when the GA DPR construct was used. We cannot rule out the possibility that GA levels derived from the G_4_C_2_ repeat transgene are below our ability to detect. As expected, neither GA nor GR were detected in animals that express the RNA-only repeat (Figure 1B). Together, these results suggest that DPR production, in particular production of the arginine-rich DPRs, drives most of the toxicity associated with the G_4_C_2_ transgene when expressed in glia. These effects are largely consistent with prior reports for expression in neurons (13,14,36–38).

### Cell-intrinsic and non-autonomous toxicity of focal induction of C9 transgenes

Having established the effects on lifespan of pan-glial expression for each of the C9orf72 transgenes, we next examined the effects of focal expression within a subset of glia in order to model the phenomenon of focal onset and intercellular propagation seen in human ALS and FTD pathology. We utilized expression in a spatially restricted glial subtype, the subperineural glia (SPG), which comprise the blood-brain barrier in *Drosophila*. These cells are advantageous for the study of focal pathological onset as they are relatively sparse in number: there are ∼300 SPG out of a total of roughly 15,000 glia in the central *Drosophila* brain (39,40). The SPG are also morphologically and spatially ideal for detecting potential non-cell autonomous effects on neurons because they are located on the surface of the brain, in a spatially distinct domain that is adjacent to the zone containing most of the neuronal somata. Thus, expression of C9orf72 products in SPG provides a spatially isolated source of each transgene’s expression, allowing examination of effects within those SPG, and in nearby neurons that lie just below the surface.

We first examined the effects on lifespan from expression of C9orf72 transgenes in the SPG, once again utilizing the Gal80^ts^ method, this time with an SPG-specific Gal4 line (SPG-Gal4, see methods). For these focal induction experiments, we selected the (G4C2)_36_ and GR_36_ transgenes, which had the most robust impact on lifespan with pan-glial expression and compared them with the RO and GA_36_ transgenes, which had more modest impacts on lifespan. As with pan-glial expression, expression of the (G_4_C_2_)_36_ and the GR DPR each caused the most robust effects on lifespan, although here, the effect with G_4_C_2_ was significantly greater than with GR. And here too, the RNA only transgene had an intermediate effect and the GA DPR had little impact on lifespan (Figure 2A). Although quantification by qPCR does reveal some modest differences in levels of expression of these transgenes (Figure S2A), these effects are not well correlated with effects on lifespan. The levels of GR DPR protein production from the (G_4_C_2_)_36_ and GR_36_ transgenes also appear roughly similar as measured by confocal microscopy (Fig. 1B).

**Figure 2.**
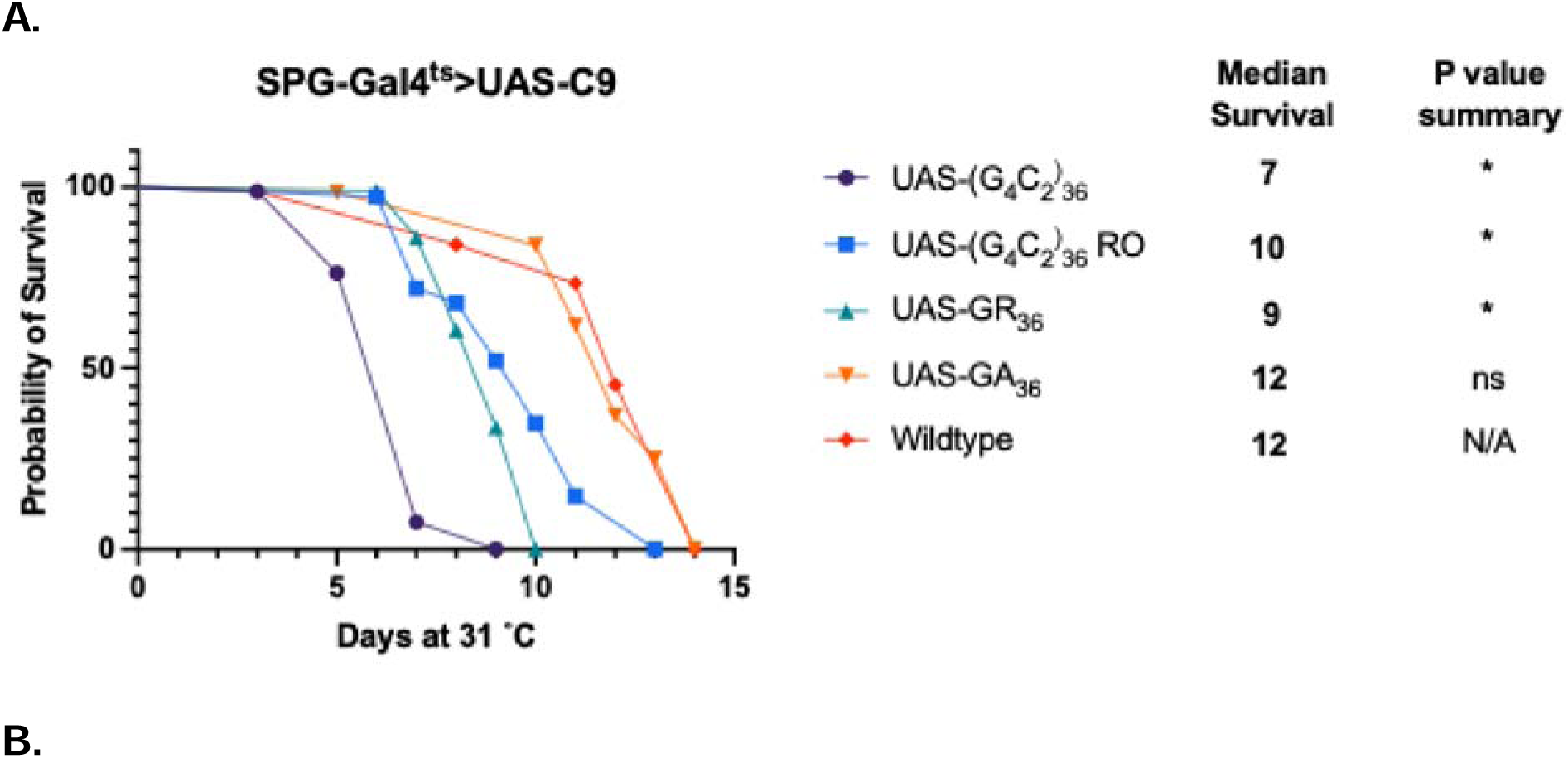
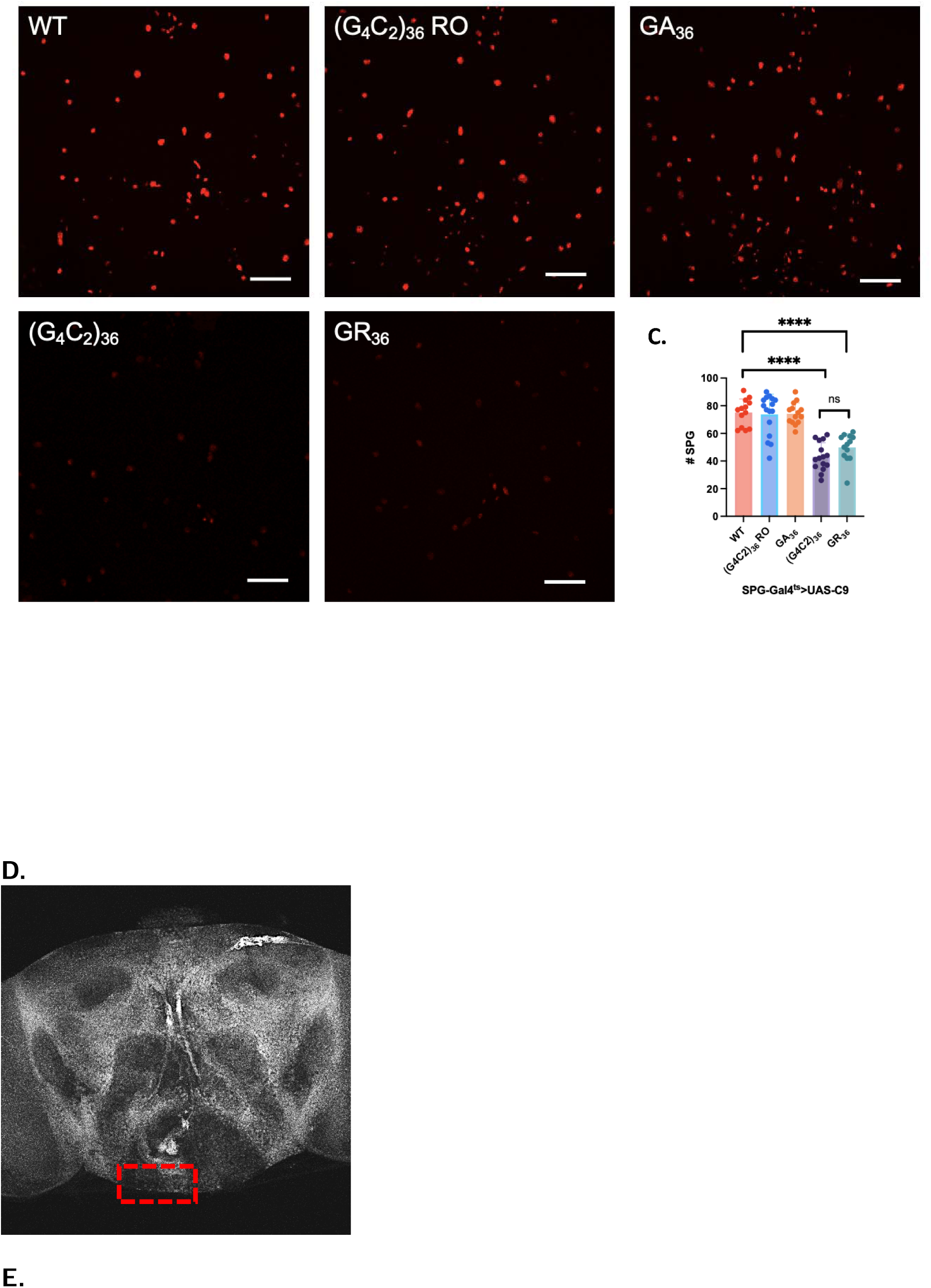
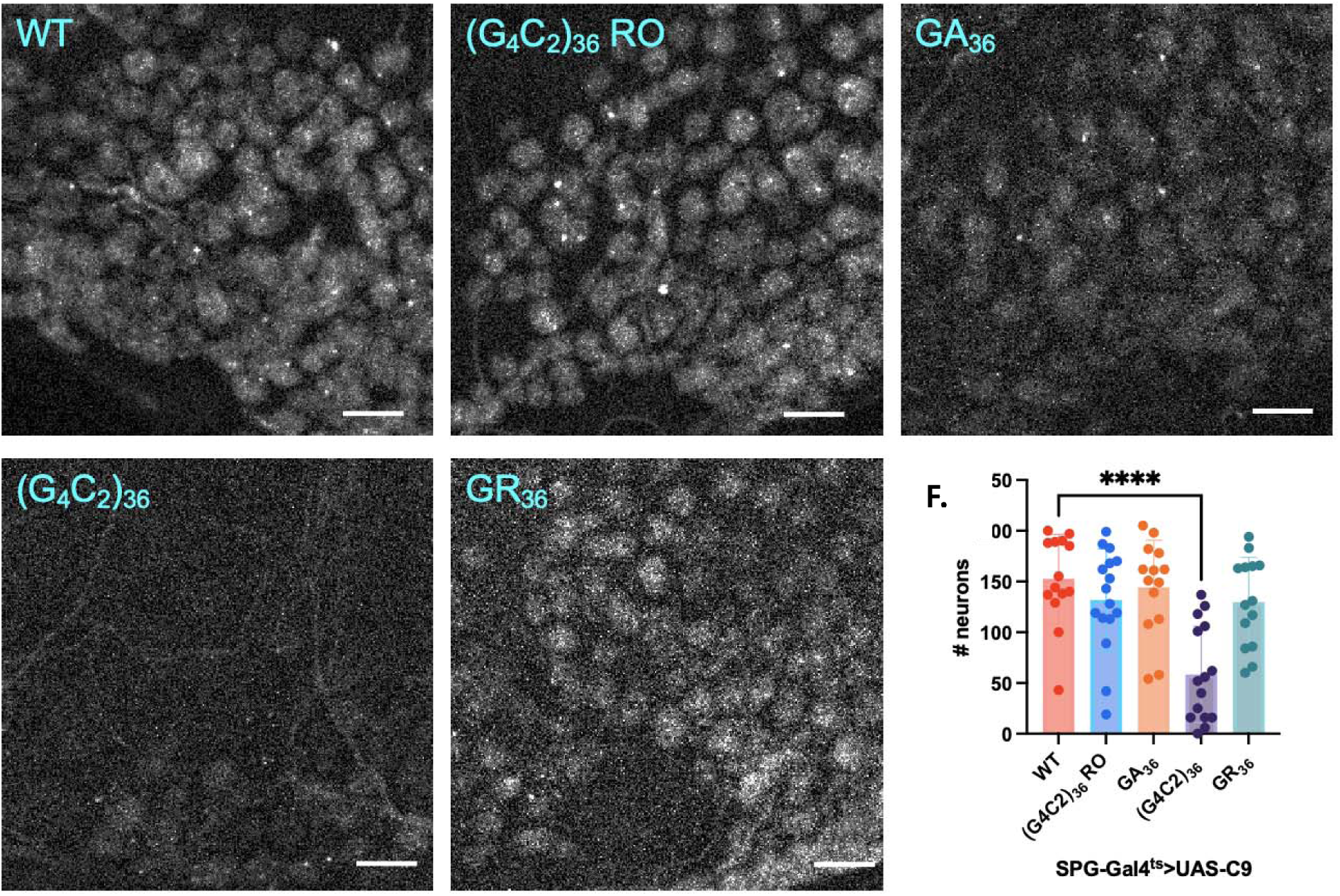
Cell-intrinsic and non-autonomous toxicity of focal induction of C9 transgenes. (A). Lifespan analysis after cell-type specific induction of C9 transgenes in supberineural glia (SPG), compared to a wildtype control with the SPG-Gal4^ts^ driver. *p < 0.005 (long-rank test with Bonferroni’s correction) (B). Z projections of confocal images through central brain five days (D5) after inducing each C9 transgene in SPG. Scale bar = 50 µM. (C) Quantification of SPG numbers at D5 after inducing C9 transgenes in SPG. ****p<0.0001 (one-way ANOVA with Dunnett’s multiple comparisons test), ns (one-way ANOVA with Šídák’s multiple comparisons test). (D). Image of *Drosophila* brain showing the location where neuronal quantification was done (red box). (E). Immunofluorescent images of neurons from region (D) of D5 brains focally expressing C9 transgenes in SPG. Scale bar = 10 µM. (F). Quantification of neurons in defined brain region of D5 brains focally expressing C9 transgenes in SPG. ****p<0.0001 (one-way ANOVA with Dunnett’s multiple comparisons test). *Full genotypes are SPG-Gal4, Tubulin-Gal80^ts^>UAS-C9 or SPG-Gal4, Tubulin-Gal80^ts^>UAS C9/UAS-WM*.

One possible explanation for the fact that the (G_4_C_2_)_36_ exhibited a more severe effect on lifespan than did the GR_36_ transgenes is that the toxicity of the (G_4_C_2_)_36_ transgene could derive from some combined production of GR plus the repeat RNA or the GA DPR, each of which are encoded by the (G_4_C_2_)_36_ transgene, but not by the GR_36_ transgene. We investigated this possibility by examining the effects of expressing combinations of both the GR_36_ and the (G_4_C_2_)_36_ RO transgenes. We also tested the combination of the GR_36_ and the GA_36_ transgenes. Each of these combinations failed to recapitulate the severity of the lifespan deficit seen with expression of just the (G_4_C_2_)_36_ transgene (Figure S2B).

To quantify cell-intrinsic effects on SPG survival, we co-expressed a nuclear H2B-mCherry reporter (UAS-WM) (41) in the SPG, along with each of the C9orf72 transgenes (Figure 2B). This reporter allowed us to label and count each of the SPG nuclei. Quantification of these images revealed that the (G_4_C_2_)_36_ and GR_36_ transgenes each caused similar reductions in numbers of H2B-mCherry labelled SPG nuclei compared to a wildtype control. By contrast, the (G_4_C_2_)_36_ RO and GA_36_ transgenes did not cause significant reduction in the numbers of SPG nuclei present (Fig. 2C). Thus, the difference in severity on lifespan effects between the (G_4_C_2_)_36_ and GR_36_ transgenes is not likely a result of differing levels of cell-intrinsic effects on survival of the SPG themselves (Figure 2C). In addition, the effects on lifespan of expressing the RNA only transgene in SPG do not appear to manifest at the level of SPG survival.

We next examined whether such focal expression in SPG of each C9orf72 repeat transgene was capable of producing non-cell autonomous effects that impacted survival of nearby neurons, as we have previously demonstrated to be the case with focal expression in SPG of a human TDP-43 expressing transgene (26). To quantify the numbers of neurons, we used immunolabeling of the pan-neuronal nuclear marker elav. For each experiment, we quantified the numbers of elav-labelled neuronal nuclei within a defined ventral region of the central brain (Figure 2D, E). Expression of the (G_4_C_2_)_36_ repeat in SPG produced a significant reduction in the number of nearby neurons. By contrast, SPG expression of either GR_36_, GA_36_, or (G_4_C_2_)_36_ RO had no significant impact on the numbers of surviving neurons (Figure 2F). These findings are consistent with the hypothesis that the reduction in lifespan that we observe in flies expressing the G_4_C_2_ repeat in SPG derives at least in part from these non-cell autonomous effects on neurons. The effects on lifespan from expressing GR_36_ may derive mostly from cell intrinsic effects on the SPG, although we cannot rule out effects on the physiology of nearby neurons.

### Expression of either (G_4_C_2_)_36_ or GR_36_ drives replication of the mdg4 endogenous retrovirus

We have previously reported that TDP-43 proteinopathy is associated with elevated expression of retrotransposons (RTEs) and endogenous retroviruses (ERVs), including the fly ERV mdg4 (29). Additionally, we previously reported that this ERV contributes to both cell-intrinsic toxicity as well as to the non-cell autonomous toxicity from glia to neurons in the *Drosophila* TDP-43 pathology model(26,27). These findings are convergent with results examining RTE expression in a TDP-43 model in mouse (42) and in postmortem cortical tissue from human subjects (30–33,43). We thus tested whether mdg4-ERV replication might also be induced in the presence of these C9orf72 transgenes. To accomplish this, we utilized the Cellular Labeling of Endogenous Retrovirus Replication of mdg4 (mdg4-CLEVR) reporter, which labels cells in which the mdg4 ERV has replicated via an RNA intermediate and then reinserted into a de novo location in the cell’s genome (41). While the replication of the mdg4-CLEVR reporter is not dependent on expression of Gal4, the expression of the fluorescent mCherry reporter after such a replication has taken place does require the presence of Gal4. In our experiments, Gal4 is only expressed in the SPG, providing the opportunity to query replication only within the SPG.

Using the CLEVR reporter, we quantified the number of SPG that exhibited mdg4 replication after 6 days of expression of each of the C9orf72 transgenes. We found that expression of either (G_4_C_2_)_36_ or GR_36_ transgenes caused significant increases in the number of SPG that exhibit mdg4 replication events, with somewhat higher numbers being observed in the GR_36_ group (Figure 3A, B). By contrast, expression of the (G_4_C_2_)_36_ RO and GA_36_ transgenes had no significant effects on mdg4 replication (Figure S3). The fact that both (G_4_C_2_)_36_ and GR_36_ caused mdg4 replication is intriguing, as only (G_4_C_2_)_36_ seems to produce measurable effects on survival of nearby neurons. This result suggests that in the context of GR-mediated toxicity, mdg4 replication in SPG may not be sufficient to cause non-cell autonomous loss of neurons as is seen with the case of TDP-43 protein pathology (26). It also suggests the possibility that some additional factor is at play with (G_4_C_2_)_36_ expression that is not present when GR_36_ is expressed.

**Figure 3.**
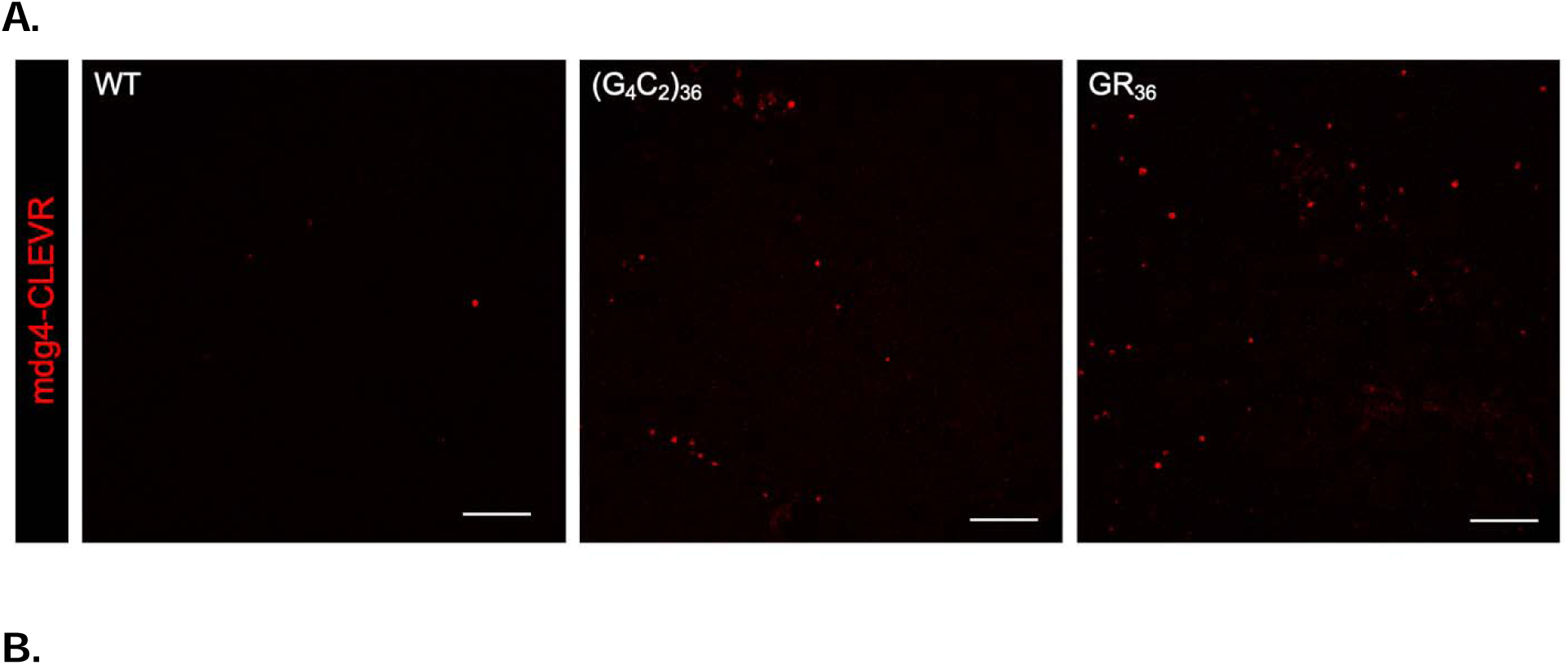
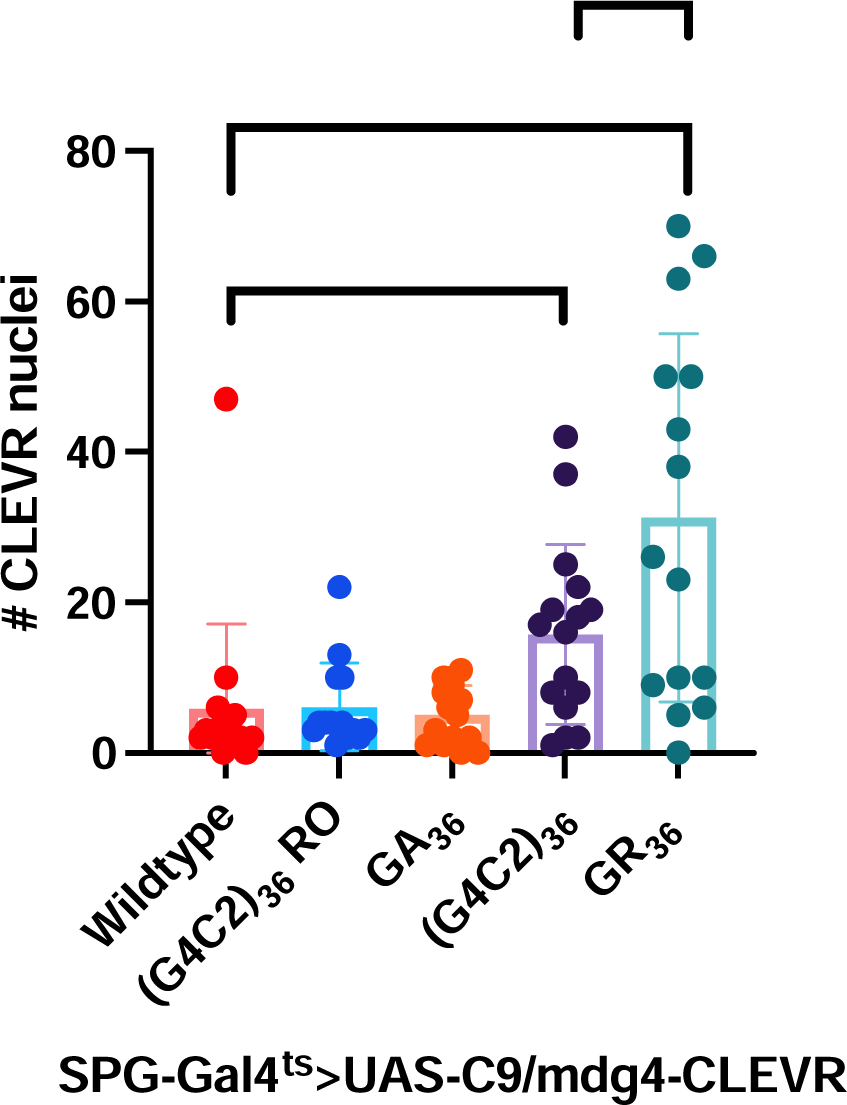
Expression of either (G_4_C_2_)_36_ or GR_36_ drives replication of the mdg4 endogenous retrovirus. (A). Confocal images showing expression of the mdg4-CLEVR reporter in anterior *Drosophila* brain at D6 of C9 transgene expression ((G_4_C_2_)_36_ RO and GA in supplement). Scale bar = 50 µm. (B). Quantification of mdg4-CLEVR reporter labelled nuclei at D6 of C9 transgene expression. *p<0.05, ***p<0.001 (one-way ANOVA with Dunnett’s multiple comparisons test), **p<0.01 (one-way ANOVA with Šídák’s multiple comparisons test). *Full genotype is SPG-Gal4, Tubulin-Gal80^ts^ > UAS-C9/mdg4-CLEVR*.

### Expression of anti- and pro-apoptosis transgenes suggests that effects on lifespan from (G_4_C_2_)_36_ and GR_36_ each involve non-cell autonomous effects

We have previously demonstrated that blocking apoptosis when TDP-43 protein pathology is induced in glia prevents the death of the glia, thereby increasing their non-cell autonomous impact to nearby neurons, and exacerbating effects on lifespan. This supports the hypothesis that the organismal toxicity produced by glial TDP-43 proteinopathy is the result of glial release of a toxic factor that impacts surrounding neurons, and that glial apoptosis may serve a neuroprotective role (26). We thus sought to examine whether a similar mechanism might be at play with expression of the toxic C9 transgenes (G_4_C_2_)_36_ and GR_36_. To test this, we compared the effects of co-expressing either an inhibitor or a promoter of apoptotic cell death. To inhibit apoptotic cell death, we used expression of p35, which directly binds to and inhibits caspase-3 (26). To promote apoptotic cell death, we co-expressed reaper (rpr) (44). We tested the effects of co-expression of p35 versus rpr in SPG along with either the (G_4_C_2_)_36_ or GR_36_ transgenes, again using the SPG-Gal4^ts^ driver, and examined the effects on lifespan. To control for the number of UAS-driven transgenes, we compared the effects of co-expressing UAS-p35 or UAS-rpr to co-expression of UAS-nls-GFP as a control (Figure 4A, 4B). We found that inhibiting apoptosis via expression of p35 extended the median lifespan of wildtype flies expressing only the SPG-Gal4^ts^ driver and no C9 transgenes by 3 days. By contrast, driving apoptosis in SPG via expression of the rpr transgene significantly shortened lifespans when this was done in animals that did not express any C9orf72 transgenes (Figure 4B). But the results were opposite when we used the same inhibitor and promoter of apoptosis in the context of (G_4_C_2_)_36_ and GR_36_ expression.

**Figure 4.**
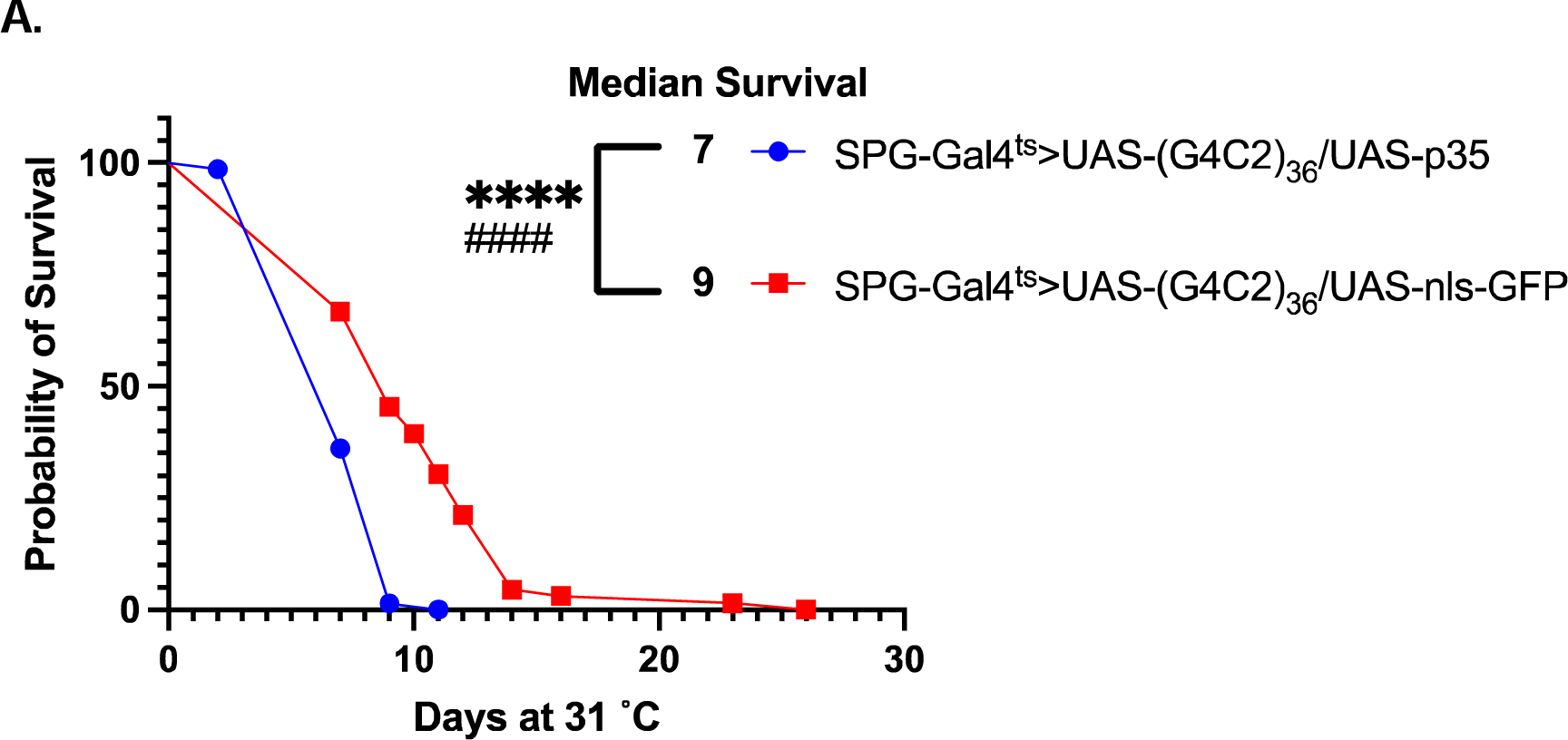
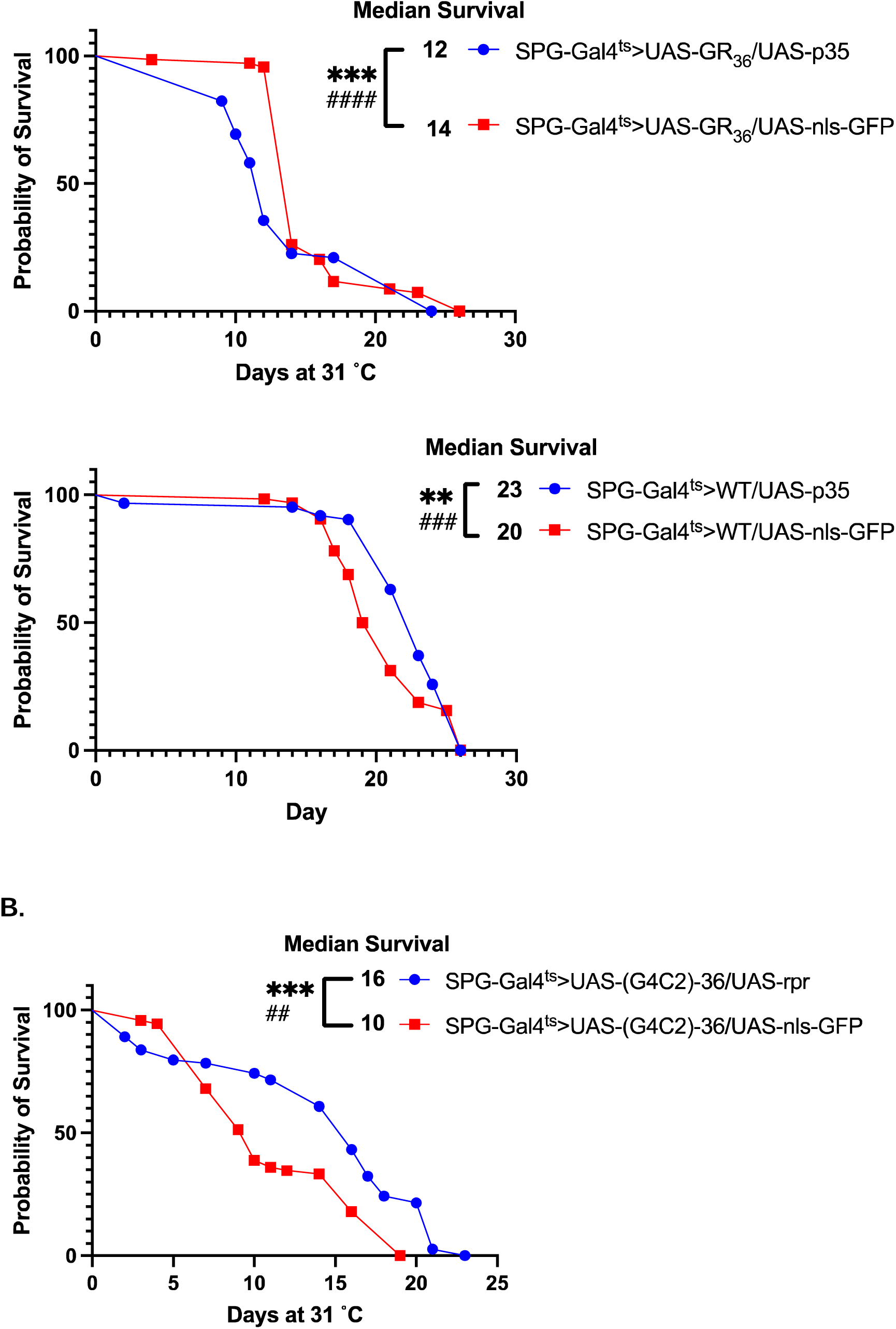
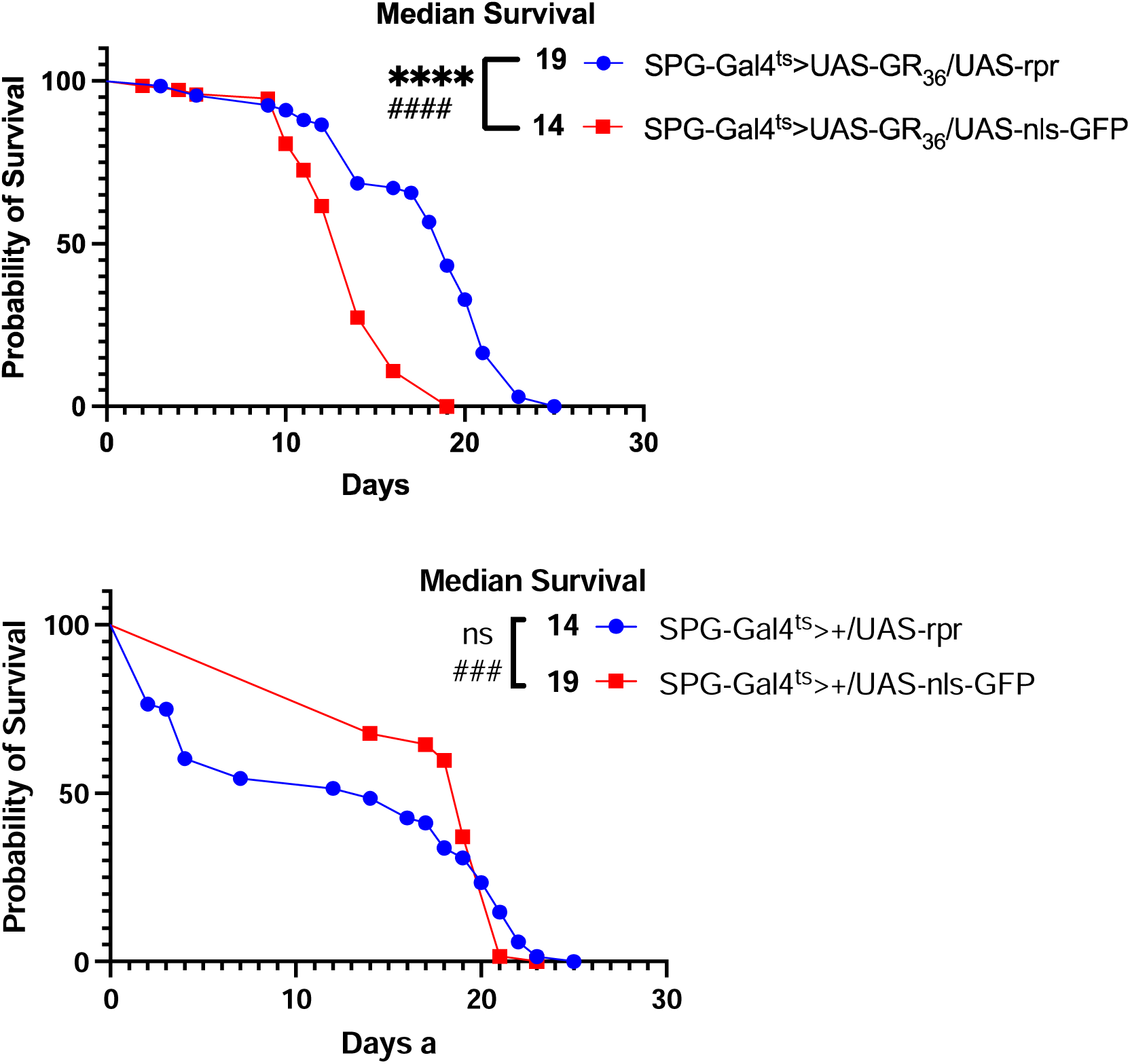
Lifespan effects of anti- and pro-apoptosis genes reveal unique modes of toxicity of G_4_C_2_ bs GR. (A). Lifespan analysis showing modifying impact of caspase inhibitor p35 on G_4_C_2_ and GR when they are co-expressed in SPG using the SPG-Gal4^ts^. Effects are compared to expression of a nls-GFP control. **p<0.01,***p<0.001, ****p<0.0001 (log-rank test), ###p<0.001, ####p<0.0001 (Gehan-Breslow-Wilcoxon test). (B). Lifespan analysis of modifying impact of expressing the pro-apoptosis gene reaper (rpr) on G_4_C_2_ and GR when expressed in SPG using the SPG-Gal4^ts^. Here too, effect is compared to expression of nls-GFP as a control. ***p<0.001, ****p<0.0001, n.s. no significant difference (log-rank test), ##p<0.01, ###p<0.001, ####p<0.0001 (Gehan-Breslow-Wilcoxon test). *Full genotypes are SPG-Gal4, Tubulin-Gal80^ts^>UAS-C9/UAS-p35 or are SPG-Gal4, Tubulin-Gal80^ts^>UAS-C9/UAS-rpr*.

When we blocked apoptosis via co-expression of p35, it further shortened the lifespans of either (G_4_C_2_)_36_ or GR_36_ expressing flies by 2 days compared to the control. This result suggests that blocking apoptosis in glial cells expressing either (G_4_C_2_)_36_ or GR_36_ exacerbated the ability of these glia to cause systemic toxic impacts to the animal’s survival. This is consistent with the idea that the SPG release a toxic substance that negatively impacts the physiology of nearby cells, and that apoptotic cell death is a means to remove these glia without release of the toxic substance. The effects with co-expressing the pro-apoptotic gene, rpr, further support this hypothesis. When we co-expressed the pro-apoptotic gene rpr along with either the (G_4_C_2_)_36_ or GR_36_ transgenes, it increased lifespan of both the (G_4_C_2_)_36_ and GR_36_ expressing flies by 6 and 5 days, respectively. This also supports the idea that apoptotic cell death of (G_4_C_2_)_36_ and GR_36_-expressing SPG is a protective mechanism.

Together, these findings with p35 and rpr are consistent with the idea that G_4_C_2_ and GR exert detrimental effects to the organism beyond their cell-intrinsic toxicity to SPG. Although we only detect non-cell autonomous effects on neuronal numbers with the (G_4_C_2_)_36_ transgene, the results of the rpr and p35 experiments are consistent with the interpretation that expression of the GR DPR may also exert some non-cell autonomous effects that impact lifespan, for example via detrimental impacts on neurophysiology.

### Blocking apoptosis in SPG increases the rate of loss of SPG glia that express (G_4_C_2_)_36_ and GR_36_

The findings described above indicate that blocking apoptosis exacerbates the non-cell autonomous effects of SPG, indicating that apoptosis of G_4_C_2_ or GR expressing glia is protective. We therefor predicted that blocking apoptosis via expression of p35 would prevent loss of the SPG glia. To test whether this was the case, we used the nuclear H2B-mCherry (UAS-WM) reporter to label SPG nuclei. This allowed us to quantify the numbers of SPG in animals that co-express (G_4_C_2_)_36_ and GR_36_ along with either p35 or an nls-lacZ as a control (to balance the number of UAS-driven transgenes across groups). Based on the findings described above for the effects of p35 on lifespan, we predicted that this manipulation would prevent loss of SPG, but cause more rapid loss of nearby neurons. But this was not the case. In contrast to what we observed with TDP-43 overexpression, we found that co-expression of p35 actually further decreased the number of SPG that survive expression of either (G_4_C_2_)_36_ or GR_36_ (Figure 5A, B). Thus paradoxically, blocking apoptosis in SPG via p35 expression leads to more substantial loss of SPG. We also quantified the non-cell autonomous effects on the number of nearby neurons and found that blocking apoptosis in SPG via expression of p35 did not cause any additional loss of neurons (Figure S5). Despite this, blocking apoptosis in these SPG does exacerbate the effects on lifespan. Taken together, these findings support the idea that in response to either (G_4_C_2_)_36_ or GR_36_ expression in SPG, cell-intrinsic effects that lead to apoptotic cell death have a protective role to the surrounding tissue and to the animal’s lifespan. When apoptotic cell death is prevented, the SPG may be caused to undergo other fates that have more detrimental systemic effects on the organism.

**Figure 5.**
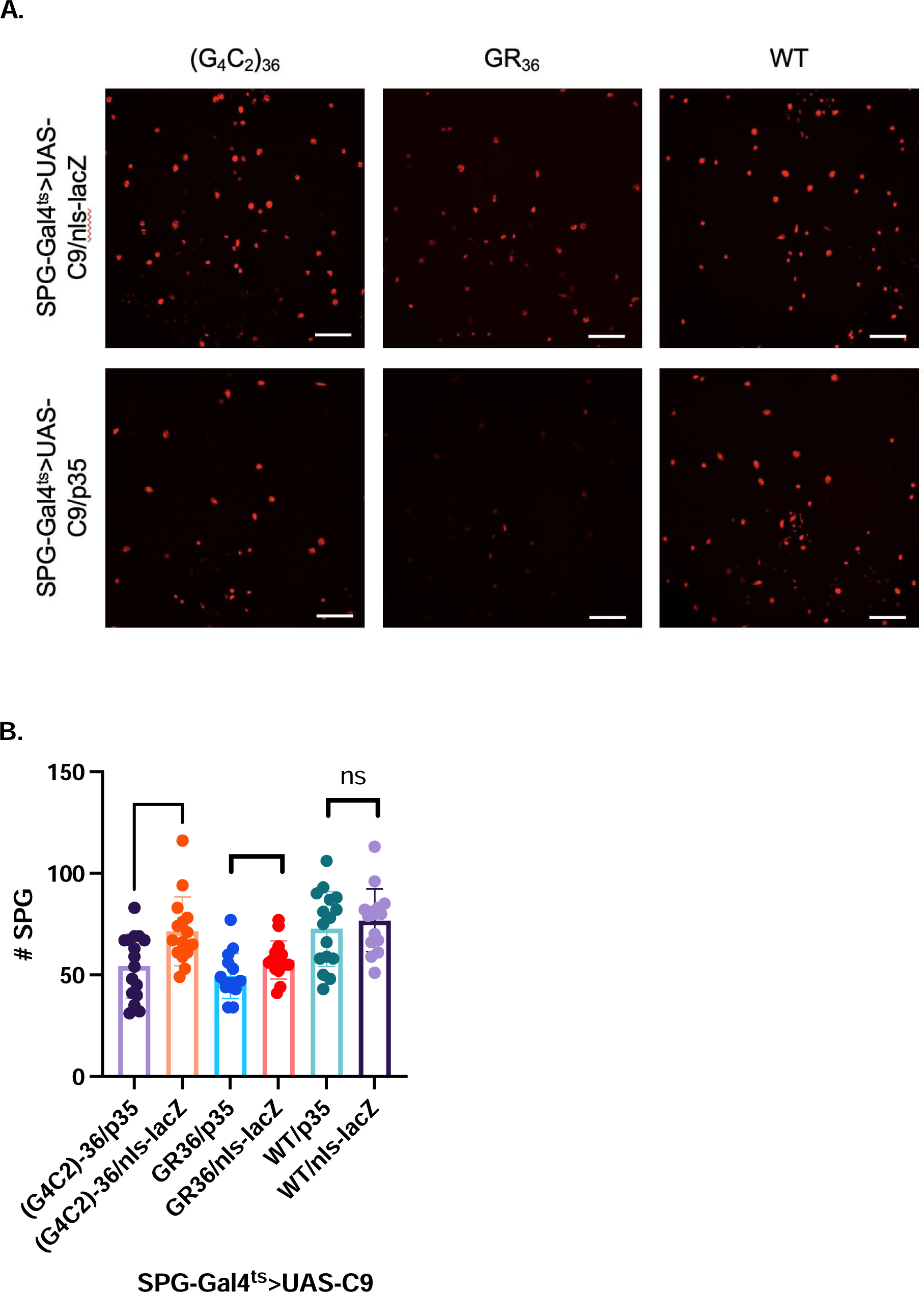
Caspase inhibitor p35 increases cell-intrinsic effects on SPG survival from (G_4_C_2_)_36_ and GR_36_. (A). Z projected confocal images through central brain of SPG after D6 of expressing C9 transgenes in SPG along with either UAS-p35 or UAS-nls-lacZ as a control. Scale bar = 50 µM. (B). Quantification of SPG numbers at D6 after induction of the C9 transgenes along with either UAS-p35 or UAS-nls-lacZ as a control. *p<0.05, **p<0.01 (Unpaired t-test). *Full genotypes are SPG-Gal4, Tubulin-Gal80^ts^>UAS-C9/UAS-p35 + UAS-WM or SPG-Gal4, Tubulin-Gal80^ts^>UAS-C9/UAS-nls-lacZ + UAS-WM*.

## Discussion

Although loss of neurons is thought to be a primary driver of clinical manifestations of neurodegenerative disorders, non-neuronal cells play a fundamental role in disease progression. Pathological changes in astrocytes, microglia, and also in several cell types that make up the blood-brain barrier (BBB) are heavily implicated in ALS, FTD, and AD (45–47). There also is evidence, both from co-culture experiments and from animal models, that astrocytes from ALS subjects become actively toxic to neurons (22,23,26,48,49). This includes the subset of cases caused by repeat expansions in C9orf72. In C9orf72 familial disease, astrocytes exhibit TDP-43 pathology (50,51) and are toxic to motor neurons grown in co-culture (24). But by contrast with TDP-43 pathology, there is little evidence from postmortem tissue for non-neuronal accumulation of C9orf72 mediated pathology itself. It is generally accepted that DPRs are not seen in postmortem glia. The RNA foci are also mostly observed in neurons, although glial cells have been reported to exhibit nuclear (but not cytoplasm) RNA foci (10,52). But observations from postmortem tissue reflect the end stage of disease, making extrapolations to what may take place during disease progression difficult. And there is clear evidence supporting the conclusion that dysfunctional physiological effects are manifested in astrocytes as well as in BBB pericytes and endothelial cells in C9orf72-driven disease (46,53,54). To date, relatively few studies have examined the *in vivo* impacts of expressing expanded C9orf72 repeats in non-neuronal cells such as glia or pericytes and endothelial cells that constitute the BBB (28,53,54). Further, there has been no systematic study of which specific components of C9orf72 gain-of-function toxicity might be responsible for producing non-cell autonomous effects on nearby neurons from non-neuronal cell types (24,25).

We use glial cell-type-specific induction of individual components of C9orf72 hexanucleotide repeat expansion pathology in the *Drosophila* brain to systematically study their relative effects on organismal lifespan, cell intrinsic toxicity, non-cell autonomous effects, and ERV activation. The transgenes capable of producing the arginine-rich sense DPR GR, namely (G_4_C_2_)_36_ and GR_36_, emerged as primary drivers of toxicity when expressed in glia, a finding in keeping with previously published results in neuronal models (13,14,36–38). To examine non-cell autonomous effects, we used focal induction within the SPG, a specialized glial cell type that constitutes the fly BBB. We used this spatially restricted expression to examine cell-intrinsic effects on survival of the SPG versus non-autonomous effects on survival of neurons. We found that compared with the GR-DPR transgene, expression in SPG of the G_4_C_2_ repeat caused more non-cell autonomous loss of neurons, despite the two lines producing relatively similar contributions to cell-intrinsic loss of the SPG. Expression of the G_4_C_2_ repeat also caused a more robust impact on lifespan than did expression of the GR-producing transgene. These results suggested several possibilities. First, production of some combination of DPRs and/or the RNA foci may both be required to mediate non-cell autonomous loss of neurons. Alternatively, the difference could lie in how GR is produced, namely by RAN translation in the G_4_C_2_ construct versus UAS-driven canonical AUG-mediated translation in the GR transgene. Since co-expressing either the RNA-only transgene or the GA_36_ transgene in combination with GR_36_ failed to recapitulate the full effects of G_4_C_2_ on lifespan, it seems unlikely that the difference has to do with combinations of factors expressed from G_4_C_2_, and rather is an effect of RAN translation from G_4_C_2_ providing the additional level of toxicity.

We have previously demonstrated that driving focal onset of TDP-43 protein pathology in *Drosophila* glia causes elevated expression of RTEs, including the fly ERV mdg4 (29). Furthermore, mdg4 activation is necessary to mediate non-cell autonomous effects that cause cell death, as well as the spread of TDP-43 pathology itself, to nearby neurons (26,27). Elevated transposable element (TE) expression has been demonstrated in C9orf72 patient brains (55,56), and C9orf72 patients exhibit TDP-43 pathology(1,57). We therefore tested whether expression of each component of C9orf72 repeat expansion pathology might recapitulate the mdg4-mediated non-cell autonomy seen in the TDP-43 model. Indeed, our results indicate that mdg4 activation is tied to expression of either the GR_36_ DPR or the G_4_C_2_ transgene, with the GR transgene exhibiting a higher degree of mdg4 replication than the G_4_C_2_ transgene. Yet, in contrast with the TDP-43 over-expression driven pathology model, mdg4 activation does not appear to be sufficient to drive non-cell autonomous loss of neurons by itself, given that only G_4_C_2_ exhibited loss of neurons in response to such SPG expression.

Another difference between the effects of expressing the G_4_C_2_ and GR transgenes compared with expression of TDP-43 is the cellular response to expression of the caspase inhibitor p35. In both the TDP-43 model (26) and the arginine dipeptide-producing C9 transgenes studied, co-expression of p35 shortened lifespan. Additionally, co-expression of the pro-apoptosis gene reaper extended lifespan in animals with SPG expression of both the G_4_C_2_ and GR transgenes. Together, these results suggest that while only expression of G_4_C_2_ in SPG produces a visible effect on the number of neurons, GR is also capable of exerting some sort of non-cell autonomous effect that impacts lifespan even though it does not result in a measurable loss of neurons. But while blocking apoptosis via p35 expression in the TDP-43 model protected glia from death and increased non-cell autonomous loss of neurons, this same manipulation in G_4_C_2_ and GR expressing SPG caused increased cell-intrinsic toxicity in glia. Although we do not fully understand the underlying mechanisms, our findings are consistent with the idea that glial expression of G_4_C_2_ and GR can lead to several different cellular outcomes. This may include more than one possible path towards programmed cell death, as well as cellular senescence (58–60). The choice of cell fate likely impacts severity and mechanisms of non-autonomous effects.

## Methods

### Drosophila strains

Flies were raised at 21 °C and maintained on standard propionic acid for all experiments described in this study. A list of transgenic lines used in this report can be found in the table below. All were backcrossed to Canton-S derivative w^118^ (*isoCJ1*), our laboratory wildtype strain, for at least five generations. Male flies were utilized throughout the study.

|  |
| --- |
| *Drosophila* Strains |
| Canton-S derivative $w^{118}$ (*isoCJ1*) |
| Repo-Gal4 |
| SPG-Gal4 |
| UAS-(G<sub>4</sub>C<sub>2</sub>)<sub>36</sub> |
| UAS-(G<sub>4</sub>C<sub>2</sub>)<sub>36</sub> RNA-only (RO) |
| UAS-GR<sub>36</sub> |
| UAS-GA<sub>36</sub> |
| UAS-PR<sub>36</sub> |
| UAS-PA<sub>36</sub> |
| UAS-WM (2<sup>nd</sup> chromosome) |
| UAS-WM (3<sup>rd</sup> chromosome) |
| mdg4-CLEVR |
| UAS-p35 |
| UAS-rpr |
| UAS-nls-lacZ |
| UAS-nls-GFP |
| tubulin-Gal80<sup>ts</sup> (3<sup>rd</sup> chromosome) |

### Lifespan analysis

All flies were incubated at 21 °C during development and shifted to a desired temperature upon eclosion. 10-15 flies of each genotype were housed within each vial, with at least 50 flies in each experimental group. Flies were counted and flipped into vials with fresh fly food every other day. The Log-Rank (Mantel-Cox) and Gehan-Breslow-Wilcoxon tests were used for survival curve comparison analysis.

### Immunohistochemistry

Adult brains were dissected in phosphate-buffered saline (PBS) at specific time points. After dissection, brains were transferred into cold 4% paraformaldyhyde (PFA) solution (1x PBS, 4% PFA, and 0.2% Triton-X 100) and fixed for two 30 minute incubation periods in a vacuum to remove trachea, with the 4% PFA replaced between fixation incubations. After fixation, dissected brains were washed three times for 10 minutes each in 1X PBST wash solution (1x PBS, 1% Triton X-100, 3% sodium chloride). Brain samples were then incubated in blocking solution (1x PBST wash solution with 10% normal horse serum) overnight at 4 °C on a nutator. After blocking, dissected brains were transferred into primary antibodies with the following dilutions: Repo (1:10, Developmental Studies Hybridoma Bank 8D12), Elav (1:10, Developmental Studies Hybridoma Bank 7E8A10), poly-GR (1:500, Sigma Aldrich ABN1361), poly-GA (1:500, Sigma Aldrich MABN889) in 1X PBST wash solution. The brains underwent primary antibody incubation overnight at 4 °C on a nutator. Brains were then washed four times for 15 minutes each in 1x PBST wash solution before being transferred to secondary antibodies (1:100) in 1x PBST wash solution with 10% normal horse solution and incubated overnight at 4 °C on a nutator. Secondary antibodies were used from the following list: donkey anti-rat DyLight 405 (Jackson ImmunoResearch Laboratories, 712-475-153), donkey anti-mouse Alexa Fluor 647 (Jackson ImmunoResearch Laboratories, 715-605-151), donkey anti-rabbit Alexa Fluor 488 (Jackson ImmunoResearch Laboratories, 711-545-152). After secondary antibody incubation, brains were washed four times for 15 minutes each in 1x PBST wash solution and then mounted on slides in FocusClear (CelExplorer).

### Confocal imaging and quantification analysis

Brain samples were imaged using a Zeiss LSM 800. All images were processed by Zeiss Zen software package and analyzed using FIJI (ImageJ). DPR staining was imaged in the central brain at 40x magnification. Watermelon (WM) and mdg4-CLEVR nuclei were imaged at 20x magnification and quantified from a Z-stack projection of the central brain. Neuronal quantification imaging was performed at 40x magnification in a standardized region of the anterior brain and quantified from a Z-stack projection.

### Statistical analysis

All statistical analyses were performed in GraphPad Prism v10.3. Unpaired t test, ordinary one-way ANOVA with multiple comparisons, Dunnett’s multiple comparisons test, Šídák’s multiple comparisons test, Log-Rank (Mantel-Cox), and Gehan-Breslow-Wilcoxon tests were used to compare the differences between groups depending on the experiment.

## Acknowledgements

We thank the Bloomington Drosophila stock center for the UAS driven C9orf72 Drosophila strains. We also wish to thank members of the Dubnau, Sher and Nelson labs for helpful discussions. This work was supported by NIA awards R01AG076493 and R01AG078788.

