## supplementary figures for "Glial cell-intrinsic and non-cell autonomous toxicity in a Drosophila C9orf72 neurodegeneration model"

**Supplemental Figures**

**A.**

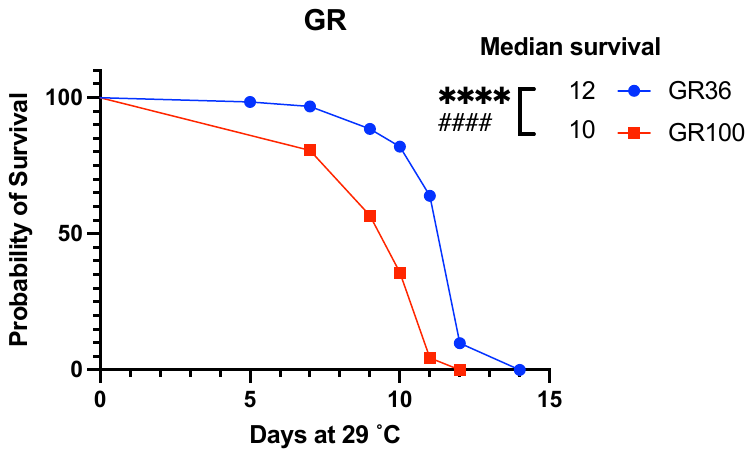

**B.**

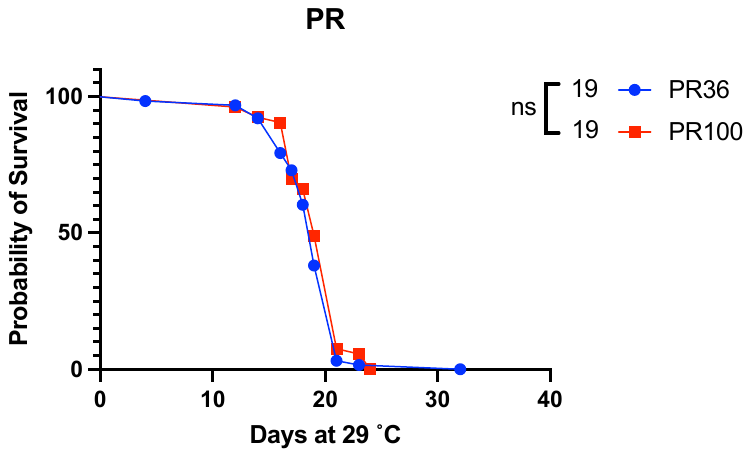

**C.**

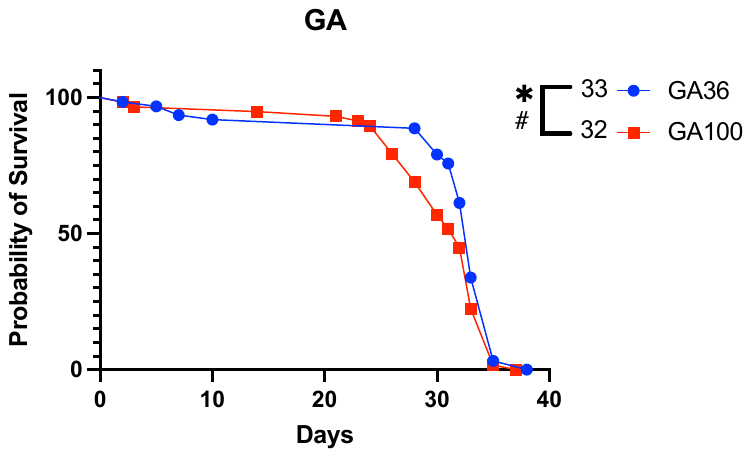

**D.**

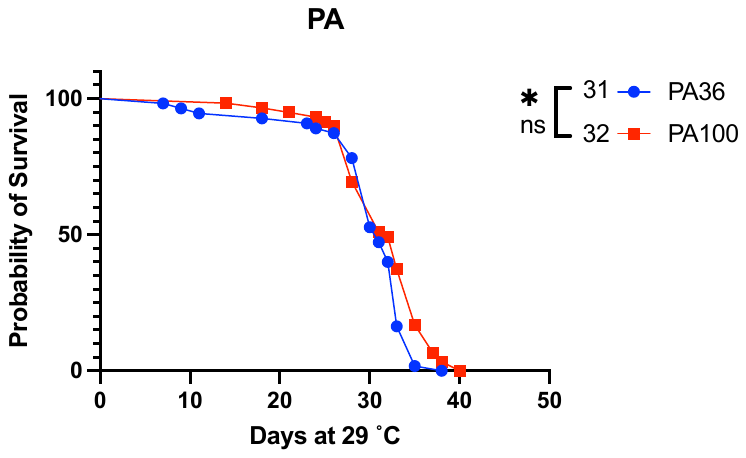

Figure S1. **Lifespan analysis of DPRs of length 36 vs. 100.** (A). GR. ****p<0.0001 (Log-Rank test), #####p<0.0001 (Gehan-Breslow-Wilcoxon test). (B). PR. ns = no significance for either Log-Rank or Gehan-Breslow-Wilcoxon test. (C). GA. *p<0.05 (Log-Rank test), #p<0.05 (Gehan-Breslow-Wilcoxon test). (D). PA. *p<005 (Log-Rank test), ns for Gehan-Breslow-Wilcoxon test. *Full genotype is Repo-Gal4, Tubulin-Gal80^ts^>UAS-C9.*

**A.**

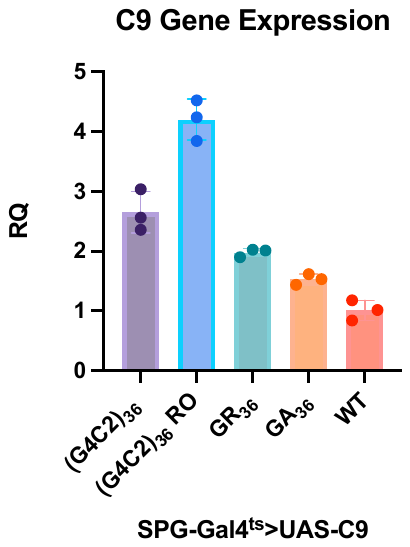

**B.**

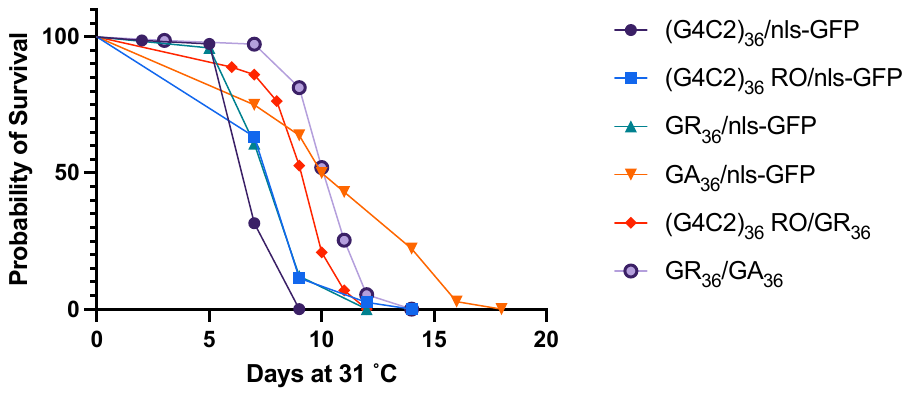

Figure S2. **Gene expression levels and combinatorial effects of C9 transgenes.** (A). Relative quantification of C9orf72 transgenes compared to actin-5C housekeeping gene. (B). Combinations of (G_4_C_2_)_36_ RO/GR_36_ and GR_36_/GA_36_ fail to recapitulate lifespan defect produced by (G_4_C_2_)_36_. *Full genotypes: SPG-Gal4, Tubulin-Gal80^ts^>UAS-C9/nls-GFP or SPG-Gal4, Tubulin-Gal80^ts^>UAS-C9_1_/UAS-C9_2_*

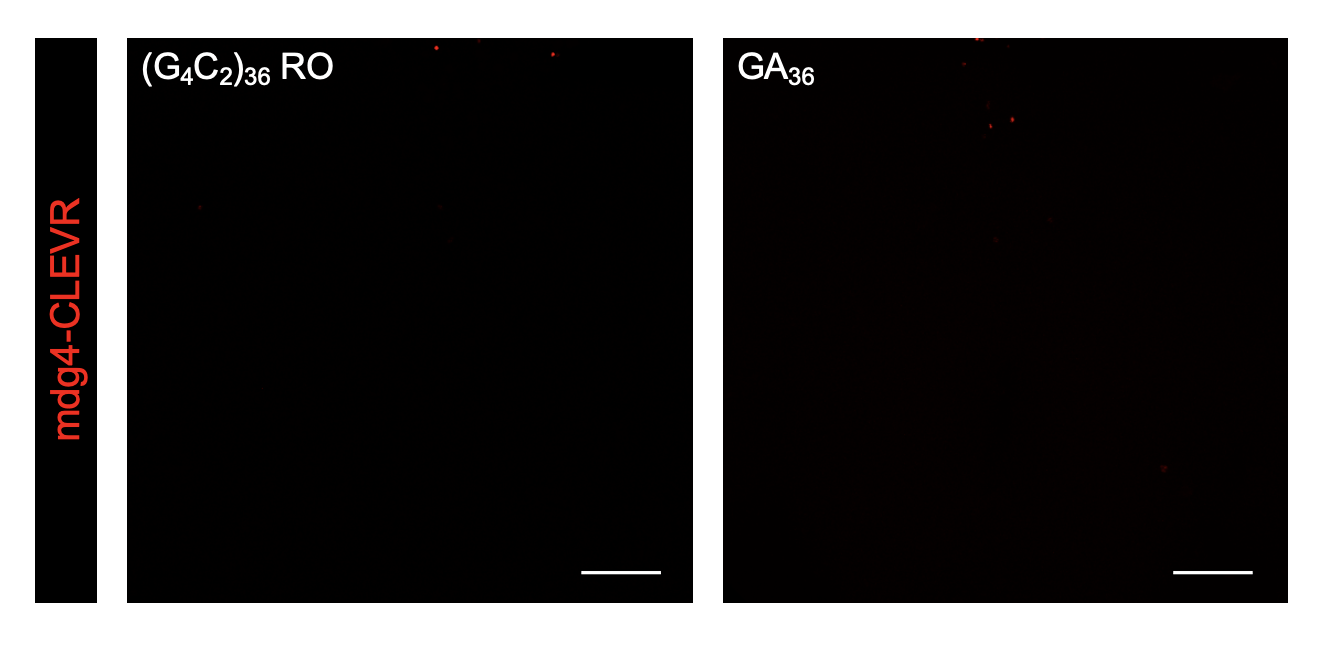

Figure S3. **mdg4-CLEVR expression in RO_36_ and GA_36_.** Scale bar = 50 µM. *Full genotypes are SPG-Gal4, Tubulin-Gal80^ts^>UAS-C9/mdg4-CLEVR.*

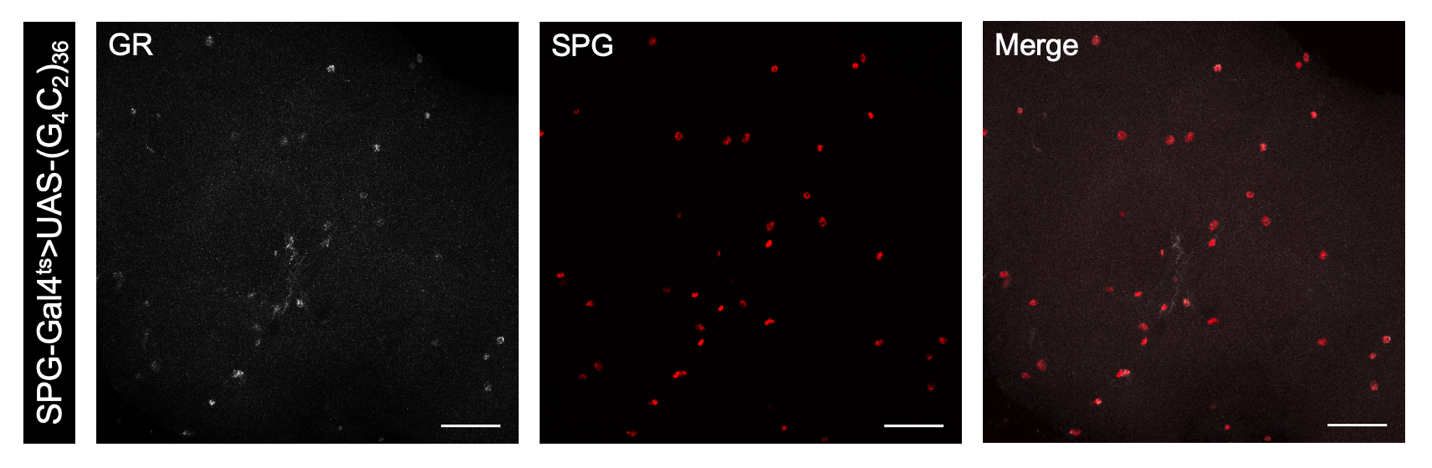

Figure S4. **GR immunolabeling in G_4_C_2_.** Scale bar = 50 µM. *Full genotype is SPG-Gal4, Tubulin-Gal80^ts^>UAS-(G_4_C_2_)_36_/UAS-WM.*

**A.**

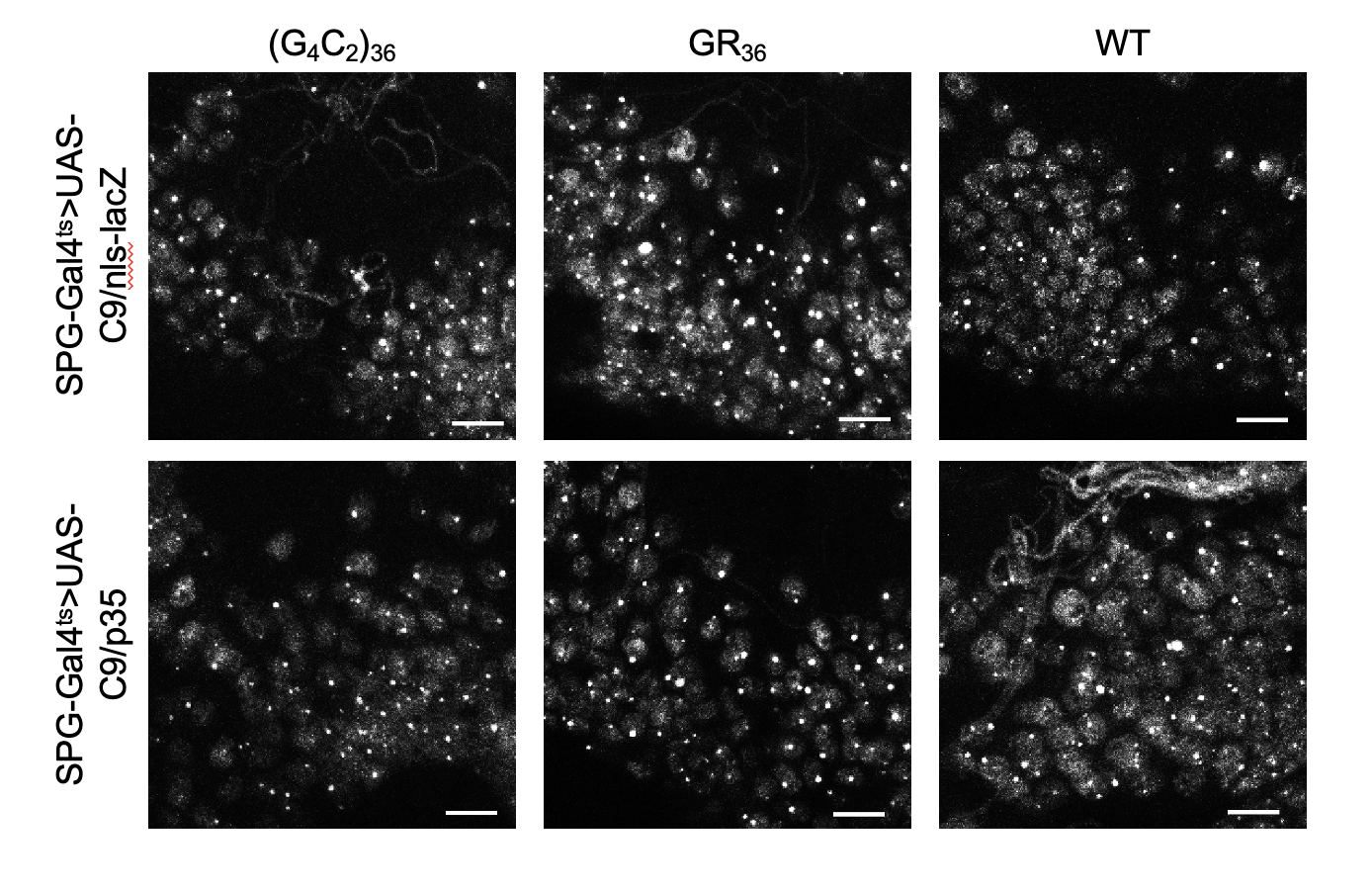

**B.**

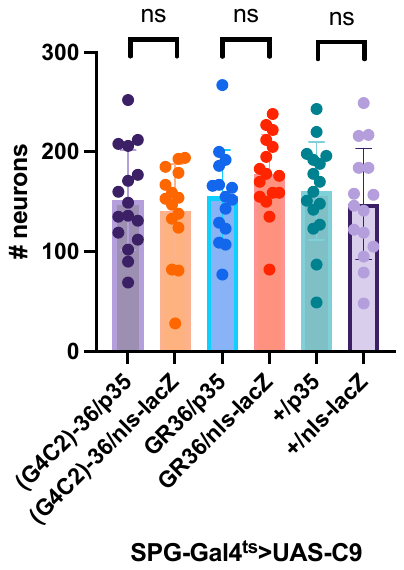

Figure S5. **Caspase inhibitor p35 does not significantly alter neuronal numbers in representative region of brain.** (A). Immunofluorescent images from neurons from D6 brains expressing C9 transgenes with either p35 or nls-lacZ control. Scale bar = 10 µM. (B). Quantification of neurons in defined region of D6 brains expressing C9 transgenes with either p35 or nls-lacZ control. ns = no significance (unpaired t-test). *Full genotypes are SPG-Gal4, Tubulin-Gal80^ts^>UAS-C9/UAS-p35 + UAS-WM or SPG-Gal4, Tubulin-Gal80^ts^>UAS-C9/UAS-nls-lacZ + UAS-WM*
